# Identification of guanylyltransferase activity in the SARS-CoV-2 RNA polymerase

**DOI:** 10.1101/2021.03.17.435913

**Authors:** Alexander P Walker, Haitian Fan, Jeremy R Keown, Jonathan M Grimes, Ervin Fodor

## Abstract

SARS-CoV-2 is a positive-sense RNA virus that is responsible for the ongoing Coronavirus Disease 2019 (COVID-19) pandemic, which continues to cause significant morbidity, mortality and economic strain. SARS-CoV-2 can cause severe respiratory disease and death in humans, highlighting the need for effective antiviral therapies. The RNA synthesis machinery of SARS-CoV-2 is an ideal drug target and consists of non-structural protein 12 (nsp12), which is directly responsible for RNA synthesis, and numerous co-factors that are involved in RNA proofreading and 5’ capping of viral mRNAs. The formation of the 5’ cap-1 structure is known to require a guanylyltransferase (GTase) as well as 5’ triphosphatase and methyltransferase activities. However, the mechanism of SARS-CoV-2 mRNA capping remains poorly understood. Here we show that the SARS-CoV-2 RNA polymerase nsp12 functions as a GTase. We characterise this GTase activity and find that the nsp12 NiRAN (nidovirus RdRP-associated nucleotidyltransferase) domain is responsible for carrying out the addition of a GTP nucleotide to the 5’ end of viral RNA via a 5’ to 5’ triphosphate linkage. We also show that remdesivir triphosphate, the active form of the antiviral drug remdesivir, inhibits the SARS-CoV-2 GTase reaction as efficiently as RNA polymerase activity. These data improve understanding of coronavirus mRNA cap synthesis and highlight a new target for novel or repurposed antiviral drugs against SARS-CoV-2.

**Importance:** SARS-CoV-2 is a respiratory RNA virus responsible for the Coronavirus Disease 2019 (COVID-19) pandemic. Coronaviruses encode an RNA polymerase which, in combination with other viral proteins, is responsible for synthesising capped viral mRNA. mRNA cap synthesis requires a guanylyltransferase enzyme; here we show that the SARS-CoV-2 guanylyltransferase is located in the viral RNA polymerase, and we identify the protein domain responsible for guanylyltransferase activity. Furthermore we demonstrate that remdesivir triphosphate, the active metabolite of remdesivir, inhibits both the guanylyltransferase and RNA polymerase functions of the SARS-CoV-2 RNA polymerase. These findings improve understanding of the coronavirus mRNA cap synthesis mechanism, in addition to highlighting a new target for the development of therapeutics to treat SARS-CoV-2 infection.

## Introduction

Coronaviruses pose a serious threat to human health as they can cause severe respiratory disease and have pandemic potential. Severe acute respiratory syndrome coronavirus (SARS-CoV) was responsible for an epidemic in 2003 which caused nearly 800 deaths, and SARS-CoV-2 is responsible for the ongoing Coronavirus Disease 2019 (COVID-19) pandemic(1). Therefore, understanding the coronavirus life cycle in order to develop novel therapeutics is of utmost importance.

SARS-CoV-2 is a betacoronavirus in the order *Nidovirales*, and it has a positive-sense RNA genome of around 30 kilobases(1, 2). The viral genome has a 5’ 7-methylguanosine (m^7^G) cap and 3’ poly(A) tail, modifications which allow the viral genome to be translated by cellular machinery. Two-thirds of the viral genome encode two overlapping open reading frames (ORFs), 1a and 1b, which are translated immediately upon infection. The resulting polyproteins are cleaved to produce non-structural proteins (nsps) 1-16, which collectively form the membrane-associated replication-transcription complex (RTC)(3–5). The RTC has several major functions: Firstly, it synthesises full-length negative-sense viral RNA, which serves as a template for new positive-sense viral RNA genomes. Secondly, it synthesises subgenomic negative-sense viral RNAs which contain the ORFs of viral structural and accessory proteins(5, 6). Finally, it transcribes the subgenomic or full-length negative-sense viral RNA to produce positive-sense, 5’ m^7^G capped, 3’ polyadenylated viral mRNAs(7, 8). Synthesising m^7^G capped viral RNA requires several distinct catalytic activities, most of which have been identified in the RTC(4). Specifically, nsp13, nsp14, and nsp16 are involved in m^7^G cap synthesis as a 5’ triphosphatase, N7-methyltransferase, and 2’-O-methyltransferase, respectively(9–12). m^7^G cap synthesis pathways also require a guanylyltransferase (GTase) enzyme, which was recently reported to reside in nsp12, the RNA-dependent RNA polymerase (RdRP) component of the RTC(13).

Like the RNA polymerases of other nidoviruses, SARS-CoV-2 nsp12 includes a 250-amino acid nidovirus RdRP-associated nucleotidyltransferase (NiRAN) domain at the N-terminus(14, 15). As the name suggests, the NiRAN domain is implicated in covalently binding to GTP and UTP nucleotides, and this domain is thought to be involved in the GTase activity of nsp12(13, 14). In order to function as a processive RNA polymerase nsp12 must form a complex with nsp7 and nsp8(16). Structures of the nsp7/8/12 RNA polymerase complex from SARS-CoV-2 have been solved by cryo-EM, and these show that nsp12 is the core of the complex while nsp8 forms an elongated scaffold around the template-product RNA duplex(17–20). Structural similarity between the nsp12 RdRP domain and other viral RNA polymerases makes the SARS-CoV-2 RNA polymerase a key target for re-purposed drugs, as nucleotide analogue drugs developed against other viruses could inhibit SARS-CoV-2 by similar mechanisms(19, 21, 22). For example, the nucleoside analogue drug remdesivir potently inhibits SARS-CoV-2 growth in cell culture and has shown potential in some clinical trials(23, 24).

Here we show that SARS-CoV-2 nsp12 has GTase activity *in vitro*, confirming the findings of another recent study(13). We further demonstrate that the NiRAN domain is responsible for this function, and we show that the active metabolite of remdesivir, remdesivir triphosphate, inhibits SARS-CoV-2 GTase activity in addition to RNA polymerase activity. These data improve understanding of the mechanism of coronavirus m^7^G cap synthesis, and highlight the NiRAN domain as a possible target for repurposed antiviral drugs.

## Results

### SARS-CoV-2 RNA polymerase purification and RNA synthesis activity

In order to gain insight into the function of the SARS-CoV-2 RNA polymerase we first aimed to establish an *in vitro* RNA synthesis activity assay, measuring extension of a 20 nucleotide (nt) primer (LS2) along a 40nt RNA template (LS1) (Fig. 1A)(16). We individually expressed wild type nsp7, nsp8 and nsp12 from SARS-CoV-2, and mixed the purified proteins to reconstitute the nsp7/8/12 RNA polymerase complex (Fig. 1B, C). In addition, we expressed and purified D760A/D761A mutant nsp12, in which two aspartic acid residues coordinating magnesium in the RdRP active site had been substituted for alanine (Fig. 1B). In the presence of RNA template and rNTPs, the wild type nsp7/8/12 complex was able to extend the LS2 primer along the LS1 template in a time-dependent manner to produce a 40nt major product (Fig. 1D). In contrast, a complex of nsp7/8/12 with D760A/D761A nsp12 was unable to extend the LS2 primer.

**FIG 1.**
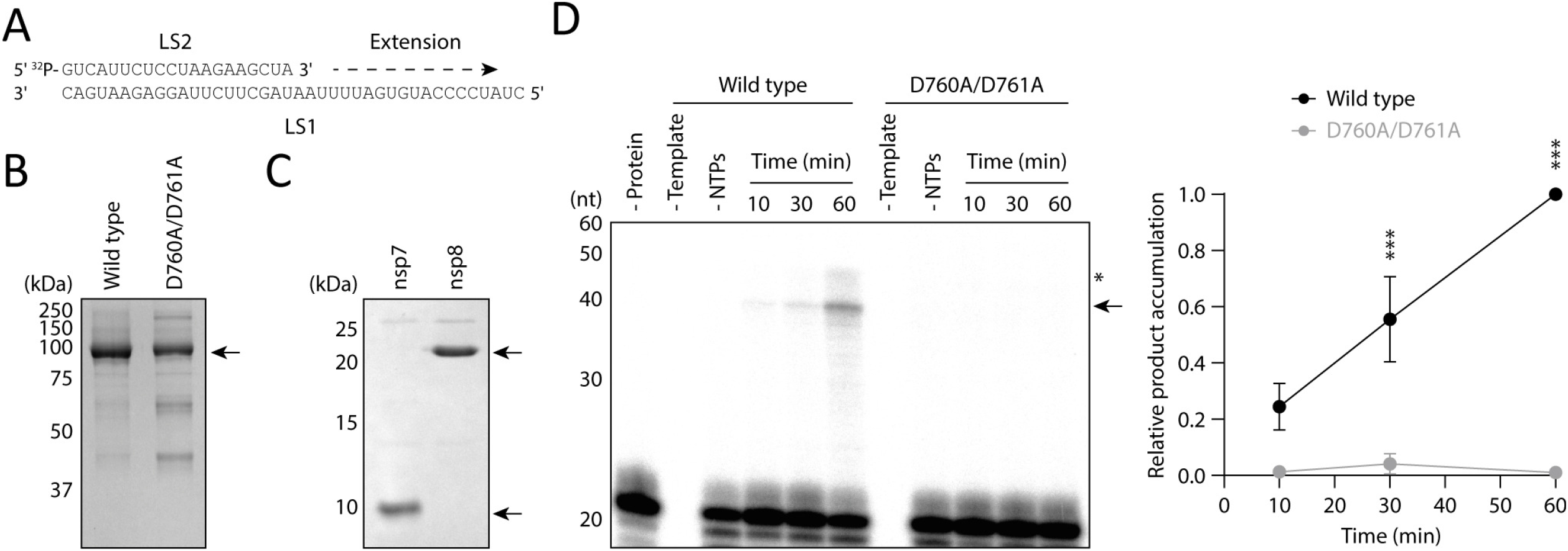
Purification and RNA polymerase activity of the SARS-CoV-2 RNA polymerase. (A) Schematic of the 40nt RNA template (LS1) and 20nt radiolabelled RNA primer (LS2). (B) Wild type and D760A/D761A mutant SARS-CoV-2 nsp12 proteins were expressed and purified from Sf9 cells and visualised by SDS PAGE. (C) SARS-CoV-2 nsp7 and nsp8 proteins were expressed and purified from E. coli cells and visualised by SDS PAGE. Arrows indicate bands corresponding to the proteins of interest. (D) Wild type or D760A/D761A mutant nsp12 was incubated with nsp7 and nsp8, then tested for *in vitro* RNA polymerase activity in the presence or absence of rNTPs and LS1/LS2 RNA template. Reaction products were resolved by denaturing PAGE and autoradiography (left). The LS2 primer and 40nt product ran slightly slower than the marker, possibly due to differences in the RNA sequence or phosphorylation state. The arrow indicates the anticipated 40nt RNA product, and the asterisk denotes incompletely denatured RNA which has slower mobility on the gel. Quantification of the 40nt RNA product is from n = 3 independent reactions (right). Data are mean ± s.e.m., analysed by two-way ANOVA. ***P<0.001.

### Nsp12 has guanylyltransferase activity

The coronavirus RTC synthesises m^7^G capped viral RNA, which requires several enzymatic activities including a GTase. GTases are responsible for covalently linking GTP to the 5’ end of diphosphorylated RNA, producing a cap-like structure (GpppN-RNA), and often involve a nucleotidylated enzyme intermediate (Fig. 2A)(25). The RNA polymerase of another nidovirus has been shown to covalently bind to GTP and UTP, which could be an intermediate step in a GTase reaction(14). Therefore, we hypothesised that a component of the SARS-CoV-2 RNA polymerase complex could be involved in m^7^G cap synthesis by functioning as a GTase.

**FIG 2.**
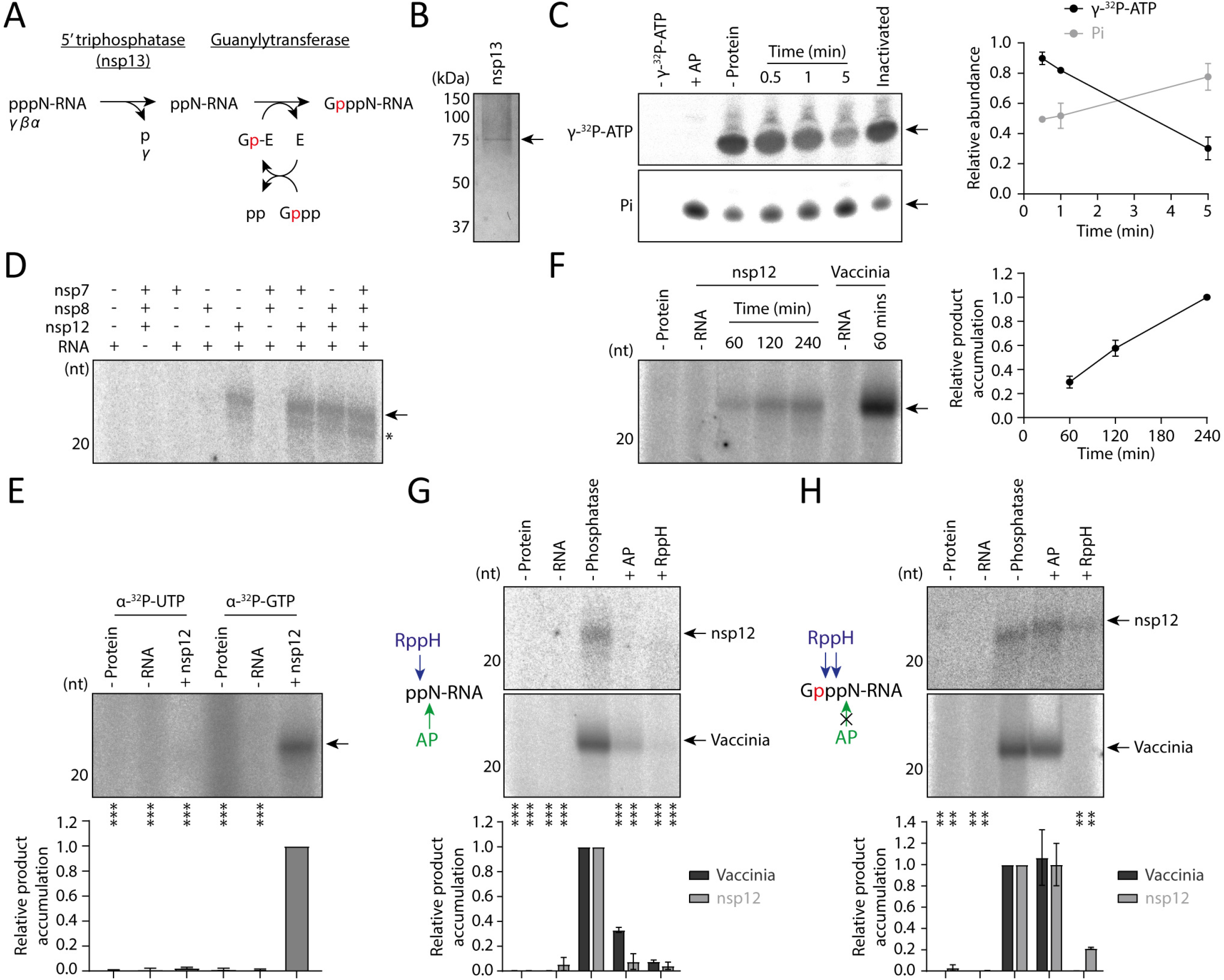
Nsp12 has guanylyltransferase activity. (A) Schematic of nsp13 5’ triphosphatase activity followed by a canonical GTase reaction mechanism, which consists of nucleotidylation of the enzyme (E) with GTP (Gppp), then transfer of GMP (Gp) to a diphosphorylated RNA substrate (ppN-RNA). The ^32^P isotope of α-^32^P-GTP is indicated in red. (B) Purified SARS-CoV-2 nsp13 protein was visualised by SDS PAGE, the arrow indicates the His6-Zbasic-tagged nsp13 band. (C) 5’ triphosphatase activity of purified nsp13 was tested by incubating with γ-^32^P-ATP for the indicated amount of time. The γ-^32^P-ATP substrate and inorganic phosphate (Pi) product were visualised by denaturing PAGE and autoradiography, arrows indicate the anticipated bands (left). Inactivated nsp13 was heated to 70°C for 5 mins prior to the reaction, and AP was used as a positive control. Quantification of γ-^32^P-ATP substrate and Pi product is from n = 2 independent reactions, data are mean ± s.e.m (right). Note that some Pi is present in the untreated γ-^32^P-ATP stock, which was subtracted from all samples as background during quantification. (D) The indicated SARS-CoV-2 RNA polymerase components were incubated with α-^32^P-GTP and diphosphorylated RNA, then radiolabelled RNA products were visualised by denaturing PAGE and autoradiography. The arrow indicates the anticipated product, and the asterisk denotes a faster mobility product which could be from contaminating RNA kinase activity in some protein preparations or decapping of the anticipated product. (E) Nsp12 was incubated with diphosphorylated RNA and α-^32^P-UTP or α-^32^P-GTP, then radiolabelled RNA products were visualised by denaturing PAGE and autoradiography. The arrow indicates the anticipated product (top). Quantification of the product is from n = 3 independent reactions (bottom). Data are mean ± s.e.m., analysed by one-way ANOVA. ***P<0.01. (F) Nsp12 or vaccinia capping enzyme were incubated with α-^32^P-GTP and diphosphorylated RNA for the indicated amount of time. Radiolabelled RNA products were visualised by denaturing PAGE and autoradiography (left), the arrow indicates the anticipated product. Quantification of the product is from n = 3 independent reactions, data are mean ± s.e.m (right). (G) Schematic of AP and RppH activity on a diphosphorylated RNA substrate (left). Diphosphorylated RNA was treated with AP or RppH, then incubated with nsp12 or vaccinia capping enzyme and α-^32^P-GTP (top). The arrow indicates the anticipated product. Quantification of the product is from n = 2 independent reactions (bottom). Data are mean ± s.e.m., analysed by one-way ANOVA. ***P<0.001. (H) Schematic of AP and RppH activity on a capped RNA substrate (left). Diphosphorylated RNA was incubated with nsp12 or vaccinia capping enzyme and α-^32^P-GTP, then reaction products were treated with AP and RppH (top). The arrow indicates the anticipated product. Quantification of the product is from n = 2 independent reactions (bottom). Data are mean ± s.e.m., analysed by one-way ANOVA. **P<0.01.

Nsp13 is thought to function directly upstream of the GTase in the m^7^G capping pathway by acting as a 5’ triphosphatase (Fig. 2A)(9). Therefore, we used purified nsp13 to generate a diphosphorylated 20nt model RNA substrate for the GTase reaction, after first confirming the *in vitro* 5’ triphosphatase activity of nsp13 on a γ-^32^P-ATP substrate (Fig. 2B, C). We incubated the diphosphorylated 20nt RNA with nsp7, nsp8, nsp12 and α-^32^P-GTP, then separated the reaction products by denaturing PAGE (Fig. 2D). In reactions which included nsp12 we observed a radiolabelled product running slightly slower than the 20nt marker, which was dependent on the presence of diphosphorylated RNA. Since the equine arteritis virus (EAV) RNA polymerase can nucleotidylate using either GTP or UTP, we also incubated nsp12 and the diphosphorylated RNA substrate with α-^32^P-UTP (Fig. 2E)(14). In this case we did not observe any radiolabelled product, suggesting that the reaction is nucleotide-specific.

To confirm that nsp12 performs a GTase reaction we compared its activity to that of vaccinia capping enzyme, a known GTase(26). When we incubated vaccinia capping enzyme with the diphosphorylated RNA and α-^32^P-GTP it produced a radiolabelled product with the same mobility as the product made by nsp12 (Fig. 2F). To further confirm that this was the capped RNA product of a GTase reaction, we performed enzymatic digestions of the substrate and product RNAs. First, we treated the substrate RNA with alkaline phosphatase (AP) or RNA 5’ pyrophosphohydrolase (RppH) to produce dephosphorylated and monophosphorylated RNAs respectively, then performed the GTase reaction (Fig. 2G). Vaccinia capping enzyme and nsp12 both displayed substantially reduced activity in the presence of dephosphorylated or monophosphorylated RNAs, indicating that a diphosphorylated RNA substrate is required. Second, after running the GTase reaction we treated the reaction products with AP or RppH (Fig. 2H). The radiolabelled products made by vaccinia capping enzyme and nsp12 were not sensitive to dephosphorylation with AP, but were sensitive to RppH. This enzymatic profile is characteristic of a GTase product; the cap-like structure in the product RNA protects 5’ phosphates from AP, however, RppH can degrade the product RNA by cleaving internally in the 5’ triphosphate(27). Collectively, these data show that SARS-CoV-2 nsp12 has GTase activity *in vitro*, which is consistent with other recent data(13).

### The nsp12 NiRAN domain is responsible for guanylyltransferase activity

Next, we investigated which domain in nsp12 is responsible for performing the GTase reaction. The EAV RNA polymerase becomes nucleotidylated in the NiRAN domain, so we hypothesised that the equivalent domain in SARS-CoV-2, consisting of nsp12 amino acid residues 1-250, could be important for GTase activity as suggested previously (Fig. 3A)(13, 14). We therefore made alanine substitutions at amino acid residues R116, D126 and D218, which are located in the NiRAN domain nucleotide binding pocket and have previously been identified as important for EAV RNA polymerase nucleotidylation activity as well as SARS-CoV growth (Fig. 3B, C)(14).

**FIG 3.**
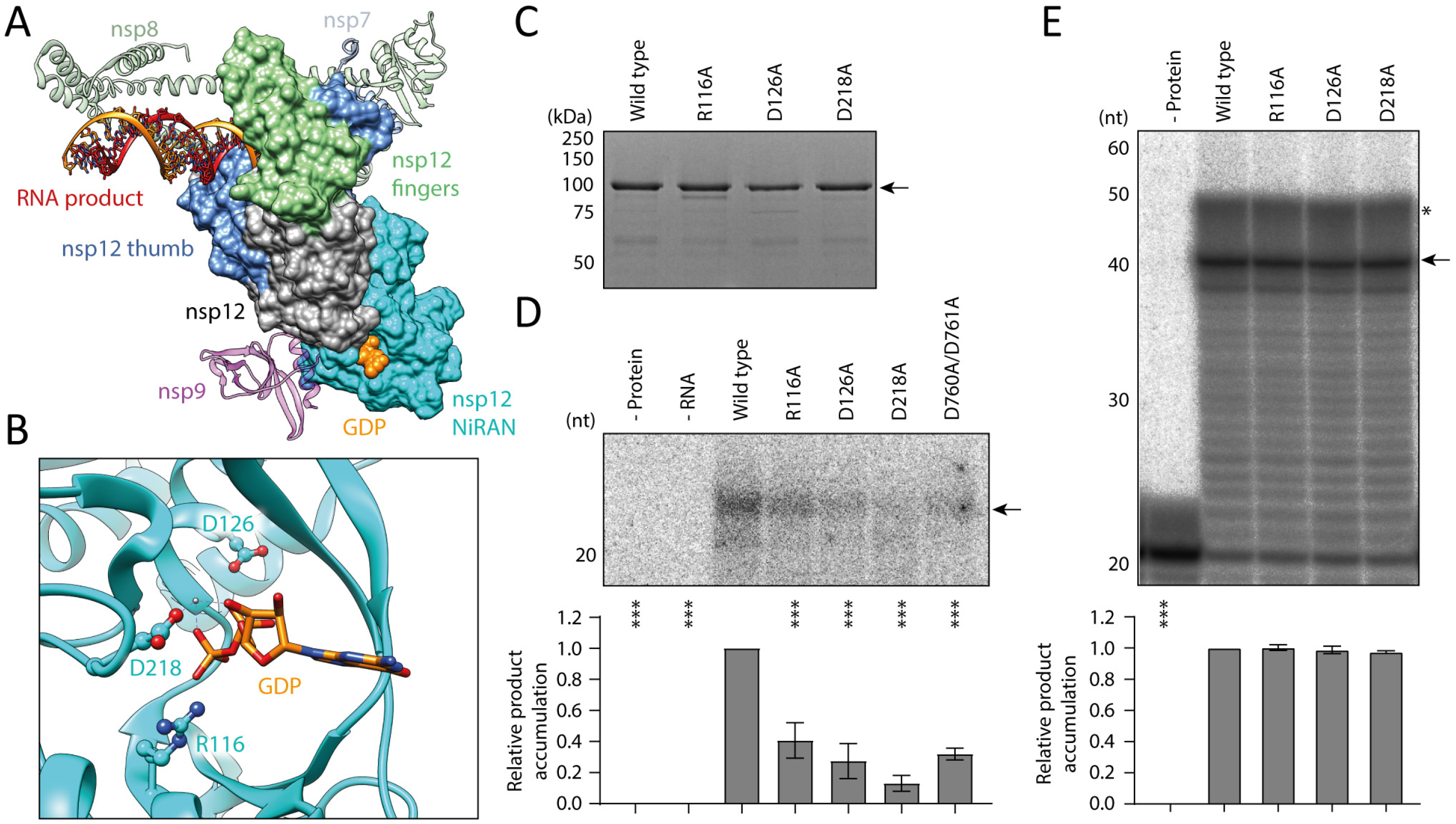
Mutations in the nsp12 NiRAN domain disrupt guanylyltransferase activity. (A) Structure of the SARS-CoV-2 nsp7/8/12 complex bound to RNA product, nsp9 and GDP (PDB: 7CYQ). Nsp12 (grey) is shown as a space-filled model with the thumb and fingers highlighted in blue and green respectively. In this structure the NiRAN domain (cyan) at the nsp12 N-terminus is bound to GDP (orange), while nsp7 and nsp8 (turquoise and green ribbons) facilitate RNA product (orange/red) binding. (B) Close-up view of GDP bound to the nsp12 NiRAN domain with key amino acid residues highlighted, including D218 which coordinates an Mg^2+^ ion (silver). (C) Mutant nsp12 proteins purified from Sf9 cells were visualised by SDS PAGE. The arrow indicates the purified nsp12 band. (D) Mutant nsp12 proteins were tested for GTase activity using α-^32^P-GTP and a diphosphorylated RNA substrate, the arrow indicates the anticipated product (top). Quantification of the product is from n = 3 independent reactions (bottom). Data are mean ± s.e.m., analysed by one-way ANOVA. ***P<0.001. (E) Mutant nsp12 proteins were tested for RNA polymerase activity in the presence of nsp7, nsp8 and LS1/LS2 RNA template (top). The arrow indicates the anticipated 40nt RNA product, and the asterisk denotes incompletely denatured RNA which has slower mobility on the gel. Quantification of the 40nt product is from n = 3 independent reactions (bottom). Data are mean ± s.e.m., analysed by one-way ANOVA. ***P<0.001.

We performed GTase reactions using the mutant nsp12 proteins, and found that all NiRAN domain mutations significantly reduced GTase activity (Fig. 3D). The D218A mutation was the most inhibitory, reducing GTase activity to approximately 10% of wild type nsp12. This result is consistent with a cryo-EM structure which shows that D218 coordinates a magnesium ion in the NiRAN domain, and therefore has a key role in nucleotide binding (Fig. 3B)(18). Interestingly, the D760A/D761A RdRP active site mutation also reduced GTase activity to approximately 40% of wild type nsp12. In contrast to their inhibitory effects on GTase activity, none of the NiRAN domain mutations significantly disrupted nsp12 RNA polymerase activity in the presence of nsp7 and nsp8 (Fig. 3E). Together, these data demonstrate that the NiRAN domain is important for GTase activity.

We then investigated whether the NiRAN domain alone is capable of performing the GTase reaction. We expressed and purified the SARS-CoV-2 NiRAN domain alone, and performed GTase reactions using either full-length nsp12 or the isolated NiRAN domain (Fig. 4A, B). The NiRAN domain was able to produce a radiolabelled product with the same mobility as the capped RNA product made by nsp12, with approximately 30% of the activity of full-length nsp12. To further confirm that the isolated NiRAN domain performs a GTase reaction, we carried out enzymatic digestions of the substrate and product RNAs. The NiRAN domain did not synthesise any radiolabelled product using dephosphorylated or monophosphorylated substrate RNA, produced by treatment of the diphosphorylated RNA with AP or RppH respectively (Fig. 4C). Furthermore, the radiolabelled product made by the NiRAN domain reaction was sensitive to degradation by RppH but not AP (Fig. 4D). These results are similar to those obtained using full-length nsp12 and vaccinia capping enzyme, confirming that the NiRAN domain alone is capable of performing the GTase reaction.

**FIG 4.**
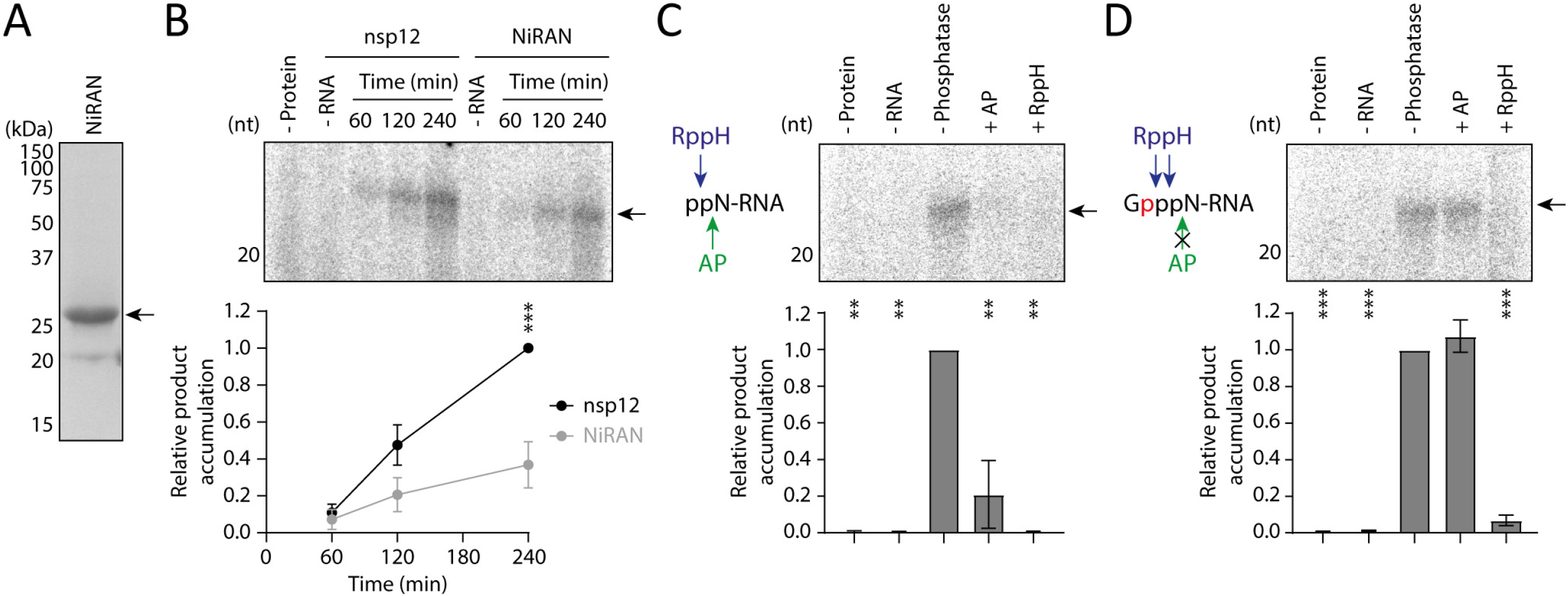
The nsp12 NiRAN domain alone has guanylyltransferase activity. (A) The NiRAN domain from SARS-CoV-2 nsp12 was expressed and purified from E. coli, then visualised by SDS PAGE. The arrow indicates the purified NiRAN domain band. (B) Full-length nsp12 and the purified NiRAN domain were tested for GTase activity using α-^32^P-GTP and a diphosphorylated RNA substrate, the arrow indicates the anticipated product (top). Quantification of the product is from n = 3 independent reactions (bottom). Data are mean ± s.e.m., analysed by two-way ANOVA. ***P<0.001. (C) Schematic of AP and RppH activity on a diphosphorylated RNA substrate (left). Diphosphorylated RNA was treated with AP or RppH, then incubated with purified NiRAN domain and α-^32^P-GTP (right). The arrow indicates the anticipated product. Quantification of the product is from n = 2 independent reactions (bottom). Data are mean ± s.e.m., analysed by one-way ANOVA. **P<0.01. (D) Schematic of AP and RppH activity on a capped RNA substrate (left). Diphosphorylated RNA was incubated with purified NiRAN domain and α-^32^P-GTP, then reaction products were treated with AP and RppH (right). The arrow indicates the anticipated product. Quantification of the product is from n = 2 independent reactions (bottom). Data are mean ± s.e.m., analysed by one-way ANOVA. ***P<0.001.

### Remdesivir triphosphate inhibits guanylyltransferase activity

Our data show that SARS-CoV-2 nsp12 has GTase activity, a function which requires nucleotide binding and therefore could be inhibited by nucleotide analogue drugs. One such nucleotide analogue is remdesivir triphosphate, active metabolite of the drug remdesivir which potently inhibits SARS-CoV-2 growth (Fig. 5A)(23).

**FIG 5.**
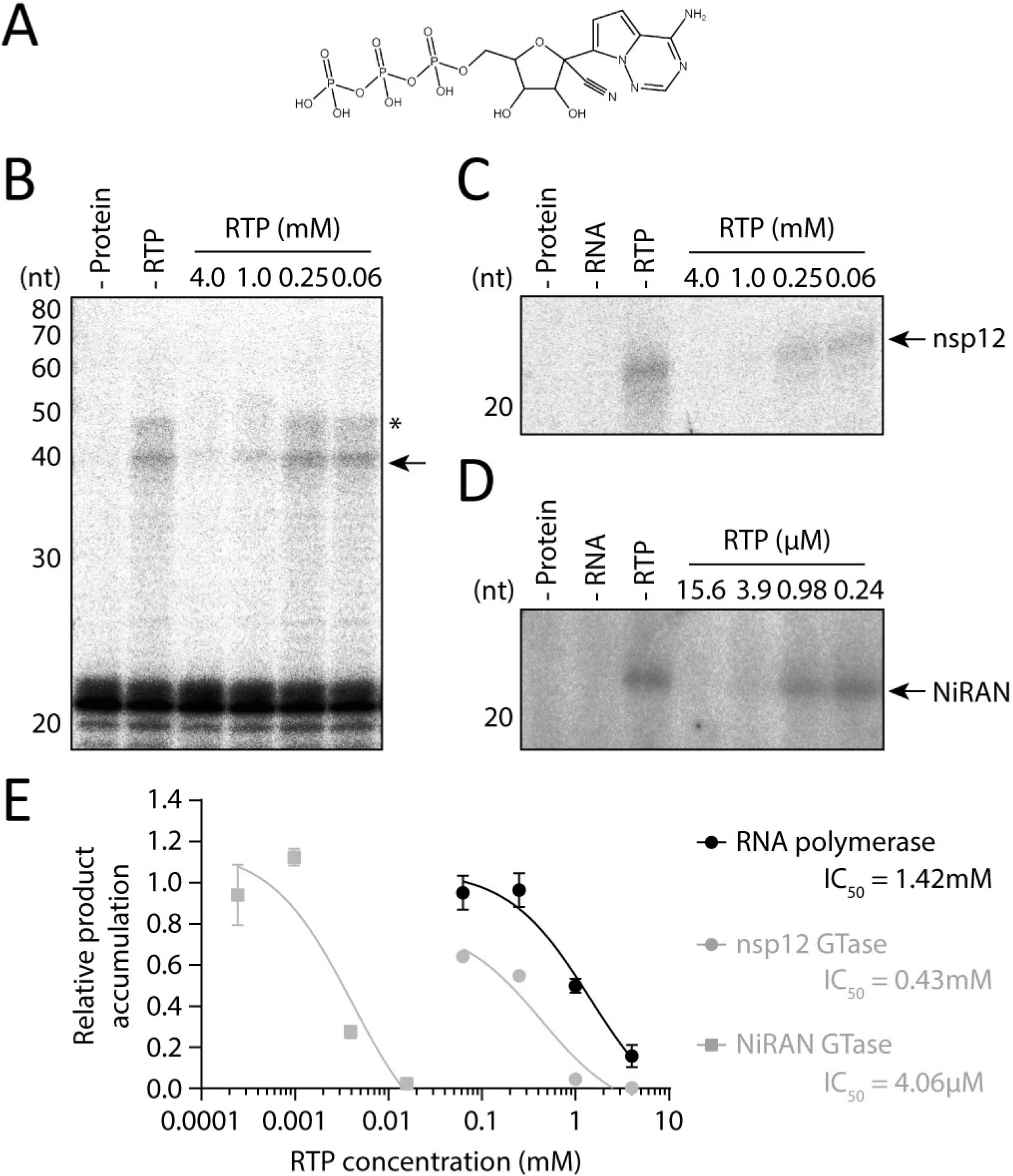
Inhibition of SARS-CoV-2 RNA polymerase and guanylyltransferase functions by remdesivir triphosphate (RTP). (A) Structure of remdesivir triphosphate, active metabolite of remdesivir. (B) Remdesivir triphosphate was titrated into RNA polymerase reactions containing nsp7/8/12 and LS1/LS2 RNA template. The arrow indicates the anticipated 40nt RNA product, and the asterisk denotes incompletely denatured RNA which has slower mobility on the gel. (C) Remdesivir triphosphate was titrated into nsp12 GTase reactions with α-^32^P-GTP and a diphosphorylated RNA substrate, the arrow indicates the anticipated product. (D) Remdesivir triphosphate was also titrated into NiRAN domain GTase reactions, the arrow indicates the anticipated product. (E) Inhibition curves for remdesivir triphosphate in SARS-CoV-2 RNA polymerase and GTase reactions. Quantification of the 40nt RNA product, nsp12 GTase product and NiRAN GTase product is from n = 3 independent reactions. Data are mean ± s.e.m., fit to dose-response inhibition curves by nonlinear regression in GraphPad Prism 9.

We first tested the inhibitory effect of remdesivir triphosphate on RNA polymerase activity using the nsp7/8/12 complex, where the drug inhibited activity with an IC_50_ of 1.42mM, which is in agreement with previous reports that remdesivir triphosphate is an RNA polymerase inhibitor *in vitro* (Fig. 5B, E)(19, 28). Next, we titrated remdesivir triphosphate into nsp12 GTase reactions, where the drug also inhibited product accumulation with an IC_50_ of 0.43mM (Fig. 5C, E). To confirm that remdesivir triphosphate inhibits the GTase reaction by binding directly to the NiRAN domain, we then titrated the drug into reactions performed using the isolated NiRAN domain (Fig. 5D, E). Under these conditions the drug also inhibited product accumulation, with a much lower IC_50_ of 4.06μM. Together, these data demonstrate that remdesivir triphosphate inhibits the GTase function of SARS-CoV-2 nsp12 in addition to RNA polymerase activity *in vitro*.

## Discussion

Despite coronaviruses being important human pathogens their mechanism of RNA synthesis remains poorly understood. Here, we show that the SARS-CoV-2 RNA polymerase has GTase activity in the NiRAN domain of nsp12.

The coronavirus RTC synthesises m^7^G capped viral RNAs, which requires several distinct catalytic activities. Most of these activities have been identified, however, until recently the viral GTase enzyme remained elusive(4, 13). With the finding that nsp12 has GTase activity, we can propose a complete model for the enzymatic processes in coronavirus m^7^G cap synthesis (Fig. 6). Nascent viral RNA is the substrate for m^7^G cap synthesis and, while the mechanism of viral RNA synthesis initiation remains unclear, reports of *de novo* initiation suggest that nascent viral RNA is 5’ triphosphorylated(16, 29–31). The γ-phosphate of nascent viral RNA is removed by the 5’ triphosphatase activity of nsp13, which interacts directly with the nsp7/8/12 complex, and the resulting diphosphorylated viral RNA can then act as a substrate for the nsp12 GTase (9, 18). Canonical GTase reactions involve a GMP-enzyme intermediate, which has been identified for the RNA polymerase of the related nidovirus EAV(14). We hypothesise that a similar SARS-CoV-2 nsp12-GMP intermediate is required for GTase activity, and the existence of this species is supported by biochemical assays(13). Following the formation of this intermediate, GMP is transferred to the diphosphorylated viral RNA to produce GpppN-RNA. This is then methylated by nsp14, which transfers a methyl group from S-adenosylmethionine (SAM) to N7 of guanine, producing S-adenosylhomocysteine (SAH) as a by-product(10, 32). Nsp16 carries out a second methylation on the 2’ hydroxyl of the nucleotide at position +1, producing viral RNA with a cap-1 structure(12, 33, 34). Higher eukaryotes have mRNA with cap-1, so this ensures that viral RNA is not recognised as non-self(35).

**FIG 6.**
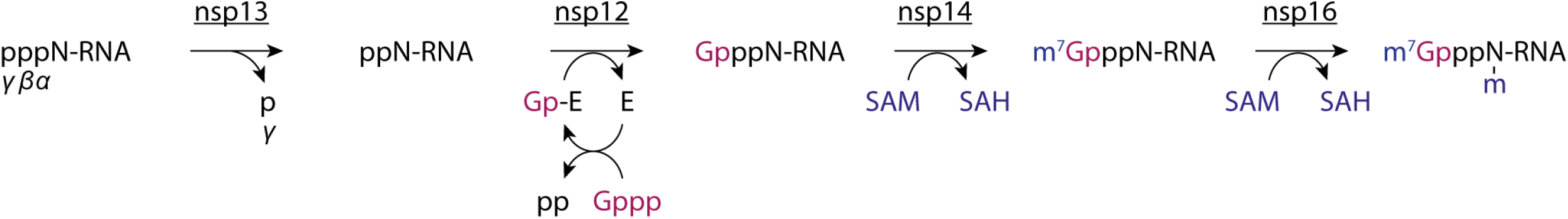
Model for coronavirus m^7^G cap synthesis. The γ-phosphate of triphosphorylated viral RNA is removed by the 5’ triphosphatase activity of nsp13. The nsp12 NiRAN domain GTase links diphosphorylated viral RNA to GTP (purple), which involves a GMP-enzyme intermediate. Nsp14 methylates GTP at the N7 position using SAM (blue) as a methyl donor, and producing SAH. Nsp16 methylates the 2’ hydroxyl group of the nucleotide at position 1 of the RNA, generating viral RNA with a cap-1 structure as the final product.

We find that the nsp12 NiRAN domain is capable of performing the GTase reaction alone, indicating this domain contains the GTase active site (Fig. 4B). However, the isolated NiRAN domain is less efficient in the GTase reaction than full-length nsp12, and mutations in the RdRP domain reduce the GTase activity of full-length nsp12 to a similar level. These results raise the possibility that the nsp12 RdRP domain is important for indirectly supporting the GTase reaction, such as by binding to the RNA substrate. The arrangement of the RdRP and NiRAN domains in nsp12 is reminiscent of non-segmented negative-strand RNA viruses (nsNSV) such as VSV, which have a GDP polyribonucleotidyltransferase (PRNTase) domain linked to the RdRP in their L proteins(36). The PRNTase has an equivalent function to the GTase in m^7^G cap synthesis, and in nsNSVs the close association of RNA synthesis and capping machinery is thought to allow viral RNA capping to take place co-transcriptionally(37). Interestingly, proteins involved in coronavirus m^7^G cap synthesis have further roles in viral RNA synthesis; specifically, nsp13 is an RNA helicase and nsp14 has a proofreading exonuclease function(9, 38, 39). This raises the possibility that coronavirus m^7^G cap synthesis also takes place co-transcriptionally, which would reduce the risk of uncapped viral RNAs activating cytoplasmic innate immune receptors such as RIG-I(35). The VSV PRNTase activity is sequence-specific, so it caps the 5’ end of viral mRNA but not the full-length viral genome(40). Our assays were performed using an RNA substrate with a sequence unrelated to the SARS-CoV-2 genome, so it remains to be seen whether the nsp12 GTase has a preference for the 5’ leader sequence shared by all SARS-CoV-2 mRNAs(8).

Viral RNA synthesis is an essential process in the SARS-CoV-2 viral life cycle, and is the target of nucleoside analogues such as remdesivir. Our *in vitro* assays indicate that remdesivir triphosphate inhibits SARS-CoV-2 RNA polymerase activity with an IC_50_ of 1.42mM, which is substantially higher than the 0.77μM IC_50_ reported for SARS-CoV-2 infections in cell culture (Fig. 5E)(23). This discrepancy could be due to the *in vitro* assay not fully reflecting the requirements for RNA polymerase activity *in vivo*; for example, in our RNA polymerase *in vitro* assay we examine the extension of an RNA primer by 20nt, whereas in an infection the RNA polymerase must processively synthesise products of up to 30 kilobases(3). Alternatively, the low IC_50_ of remdesivir *in vivo* could be explained by a dual mechanism of action. We find that remdesivir triphosphate inhibits nsp12 GTase activity with a slightly lower IC_50_ than RNA polymerase activity *in vitro*, and it is unclear which of these inhibitory activities is most important for reducing viral growth (Fig. 5E). We also find that the drug inhibits the NiRAN domain GTase reaction with a much lower IC_50_, which could be due to mechanistic differences in the GTase reaction compared to full-length nsp12, or because remdesivir triphosphate binding to the nsp12 RdRP domain reduces efficacy. The SARS-CoV-2 nsp12 NiRAN domain is an attractive target for future antiviral drugs as it could be targeted by nucleotide analogues, and *in silico* studies propose that certain kinase inhibitors may be able to bind to the NiRAN domain(41). The GTase activity assay we have established here could function as a useful tool to assess the potency of these compounds.

In summary, we have identified that the SARS-CoV-2 RNA polymerase is a GTase enzyme, which provides a complete enzymatic model for coronavirus m^7^G cap synthesis and is consistent with a recent independent study(13). We additionally show that the NiRAN domain is responsible for GTase activity, which presents a new target for novel or repurposed antiviral drugs against SARS-CoV-2.

## Materials and methods

### SARS-CoV-2 protein expression and purification

Full-length nsp7, nsp8 and nsp12 genes from SARS-CoV-2, codon optimized for insect cells, were purchased from IDT. The nsp12 gene was cloned into the MultiBac system, with a Tobacco Etch Virus (TEV) protease cleavable protein-A tag on the C-terminus(42). Nsp7, nsp8 and the NiRAN construct (nsp12 amino acid residues 1-259) were cloned into pGEX-6P-1 vector (GE Healthcare) with an N-terminal GST tag followed by a PreScission protease site. Nsp12 was expressed in Sf9 insect cells and nsp7, nsp8 and NiRAN were expressed in E. coli BL21 (DE3) cells. Initial purification of nsp12 was performed by affinity chromatography as previously described for the influenza virus RNA polymerase with minor modifications: all buffers were supplemented with 0.1mM MgCl_2_ and the NaCl concentration was changed to 300mM(43).

Nsp7, nsp8 and NiRAN were purified on Glutathione Sepharose (GE Healthcare). After overnight cleavage with TEV (nsp12) or PreScission (nsp7, nsp8 and NiRAN) proteases, the released proteins were further purified on a Superdex 200 (for nsp12) or a Superdex 75 (for nsp7, nsp8 and NiRAN) Increase 10/300 GL column (GE Healthcare) using 25mM HEPES–NaOH, pH 7.5, 300mM NaCl and 0.1 mM MgCl_2_. Fractions of target proteins were pooled, concentrated, and stored at 4°C.

### RNA polymerase *in vitro* activity assays

Nsp7, nsp8 and nsp12 were mixed at a molar ratio of 5:5:1 and incubated on ice for 1 hour to form the nsp7/8/12 complex. Activity assays were performed essentially as described previously for SARS-CoV nsp7/8/12(16). Briefly, 40mer LS1 (5’-CUAUCCCCAUGUGAUUUUAAUAGCUUCUUAGGAGAAUGAC-3’) and radiolabelled 20mer LS2 (5’-GUCAUUCUCCUAAGAAGCUA-3’) RNAs corresponding to the 3’ end of the SARS-CoV genome (without the polyA tail) were pre-annealed by heating to 70°C for 5 mins followed by cooling to room temperature. 50nM pre-annealed RNA was incubated for the indicated time at 30°C with 500nM nsp7/8/12 complex, in reaction buffer containing 5mM MgCl_2_, 0.5mM of each ATP, UTP, GTP and CTP, 10mM KCl, 1U RNasin (Promega) and 1mM dithiothreitol (DTT), and remdesivir triphosphate or ribavirin triphosphate was included where indicated. Reactions were stopped by addition of 80% formamide and 10mM EDTA, followed by heating to 95°C for 3 mins. Reaction products were resolved by 20% denaturing PAGE with 7M urea, and visualised by phosphorimaging on a Fuji FLA-5000 or Typhoon FLA-9500 scanner. Data were analysed using ImageJ and Prism 9 (GraphPad).

### Generation of diphosphorylated RNA using nsp13

Purified SARS-CoV-2 nsp13 with an N-terminal His6-Zbasic tag in 25mM HEPES–NaOH, pH 7.5, 300mM NaCl and 5% glycerol was a kind gift from Yuliana Yosaatmadja and Opher Gileadi. A 5μM 1:1 mixture of di- and triphosphorylated 20mer model RNA (5’-AAUCUAUAAUAGCAUUAUCC-3’) (Chemgenes) was treated with 250nM SARS-CoV-2 nsp13 in the presence of 5mM MgCl_2_ for 5 mins at 30°C, followed by heat inactivation of nsp13 at 70°C for 5 mins. The resulting diphosphorylated RNA stock was used for all GTase reactions. To test nsp13 activity, 4.75μM ATP mixed with 0.25μM γ-^32^P-ATP was used as a substrate. As a positive control, 0.5U/μl FastAP Thermosensitive Alkaline Phosphatase (Thermo) was incubated with 4.75μM ATP and 0.25μM γ-^32^P-ATP for 1 hour at 37°C. Reactions were stopped by addition of 80% formamide and 10mM EDTA, followed by heating to 95°C for 3 mins. Reaction products were resolved by 20% denaturing PAGE with 7M urea, and visualised by phosphorimaging on a Fuji FLA-5000 scanner. Data were analysed using ImageJ and Prism 9 (GraphPad).

### Guanylyltransferase *in vitro* activity assays

Complexes of nsp7, nsp8 and nsp12 were formed as described above, and nsp12 was used for GTase reactions unless indicated otherwise. Where indicated, 5μM diphosphorylated 20mer RNA substrate (see above) was pre-treated with 0.3U/μl FastAP Thermosensitive Alkaline Phosphatase (Thermo) at 37°C for 1 hour. Alternatively, 5μM diphosphorylated 20mer RNA substrate was pre-treated with 0.5U/μl RNA 5’ Pyrophosphohydrolase (NEB) in 1× NEBuffer 2 (NEB) at 37°C for 1 hour. Enzymes were heat inactivated at 75°C for 5 mins before the resulting RNA substrates were used in GTase reactions.

To run the GTase reaction, 500nM protein was incubated with 1μM diphosphorylated or phosphatase-treated 20mer RNA, 0.05μM α-^32^P-UTP or α-^32^P-GTP, 5mM MgCl_2_, 10mM KCl, 1U RNasin (Promega) and 1mM DTT at 30°C for 240 mins unless stated otherwise. GTase reactions involving vaccinia capping enzyme were run for 60 mins under the same conditions, using 0.01U/μl vaccinia capping enzyme (NEB) instead of SARS-CoV-2 protein.

Where indicated, completed GTase reactions were treated with 0.3U/μl FastAP Thermosensitive Alkaline Phosphatase (Thermo) at 37°C for 1 hour. Alternatively, the reactions were treated with 0.5U/μl RNA 5’ Pyrophosphohydrolase (NEB) in 1x NEBuffer 2 (NEB) at 37°C for 1 hour. All reactions were stopped by addition of 80% formamide and 10mM EDTA, followed by heating to 95°C for 3 mins. Reaction products were resolved by 20% denaturing PAGE with 7M urea, and visualised by phosphorimaging on a Fuji FLA-5000 or Typhoon FLA-9500 scanner. Data were analysed using ImageJ and Prism 9 (GraphPad).

## Acknowledgements

We thank Yuliana Yosaatmadja and Opher Gileadi for providing purified SARS-CoV-2 nsp13.

This work was supported by Medical Research Council (MRC) programme grant MR/R009945/1 to E.F., Wellcome Investigator Award 200835/Z/16/Z to J.M.G. and MRC Studentship to A.P.W. This work was also supported by the COVID-19 Research Response Fund administered through the Medical Sciences Division of the University of Oxford.

All authors have contributed to the design of the experiments. A.P.W., H.F. and J.R.K. performed the experiments and A.P.W. analysed the data. A.P.W. and E.F. wrote the paper with input from all authors.

We have no conflicts of interest to declare.

